# Low-Frequency Textured Gabor Flicker Enhances Neural Entrainment and Visual Comfort for Brain-Computer Interface Control

**DOI:** 10.64898/2025.12.12.693875

**Authors:** Jules Gomel, Marie-Constance Corsi, Frédéric Dehais

## Abstract

Steady-state visual evoked potentials (SSVEPs) are widely used in cognitive neuroscience and brain–computer interfaces (BCIs), but the visual discomfort induced by repetitive luminance flicker limits their usability, particularly in multi-target settings due to strong peripheral distraction. Textured flicker composed of Gabor patches has been proposed as a more comfortable alternative, but its suitability for SSVEP paradigms and its frequency-dependent impact on neural entrainment remain unclear. Here, we directly compared textured Gabor-based flicker and classical luminance flicker using a frequency sweep followed by a multi-class SSVEP BCI task. In Session 1 (N=24), we measured SSVEP signal-to-noise ratio (SNR), inter-trial coherence (ITC), and subjective comfort across 13 stimulation frequencies (3–18Hz). Gabor-based textures elicited higher SNR and ITC than plain flicker at low frequencies (3–9Hz), whereas plain flicker produced stronger and more phase-consistent responses at higher frequencies (12–18Hz), revealing a robust crossover in entrainment. Across almost all frequencies, Gabor stimuli were rated as more comfortable. Based on these results, we defined a low-frequency Gabor-optimized band (5–7Hz) and a higher plain-optimized band (14–16Hz). In Session 2 (N=18), these bands were used in a five-class offline SSVEP BCI. Classification accuracy was highest (Gabor: 95.7% at 5–7Hz; plain: 98.1% at 14–16Hz) when each stimulus type was used in its optimal band and decreased markedly when stimulus type and frequency band were mismatched. Gabor stimuli were consistently rated as more comfortable and nearly imperceptible in peripheral vision. Together, these findings establish textured Gabor flicker as a comfortable and effective alternative to luminance flicker for low-frequency SSVEP paradigms.

## 1 Introduction

Rhythmic sensory input can phase-align ongoing brain activity, tuning endogenous oscillations to the temporal structure of external events (Lakatos et al., 2019). In vision, such alignment produces steady-state visual evoked potentials (SSVEPs): sustained, frequency-locked responses—maximal over occipital cortex—elicited by rapid visual stimuli (RVS) and robustly captured with electroencephalography (EEG). Because of their reliability and spectral precision, SSVEP paradigms are widely used in cognitive neuroscience to probe cognition (Norcia et al., 2015). Frequency-tagging paradigms, which present RVS at predefined rates, afford fine-grained tests of oscillation-cognition links—including attention (Gulbinaite et al., 2019), face perception (Boremanse et al., 2013), working memory (Ellis et al., 2006), and language (Wang et al., 2021)—and enable rhythm-specific visual entrainment (Gulbinaite et al., 2017; Schwab et al., 2006; Wiesman et al., 2019).

In Brain-Computer Interfaces (BCIs), which aim to translate brain activity into commands (McFarland and Wolpaw, 2011), frequency coding yields many selectable classes with distinctive spectral features, supporting high information-transfer rates (ITR) and strong accuracy (F.-B. Vialatte et al., 2010). Such performance levels are made possible by advanced decoding techniques—most notably Task-Related Components Analysis (TRCA)—which extract highly reproducible SSVEP components and support the high discriminability required for large-scale frequency-coded BCIs, such as a 40-target keyboard with ITRs above 200 bit/min (Nakanishi et al., 2018). Despite these strengths, flicker-based stimulation carries important limitations. Increasing luminance or contrast can boost response amplitude, but sustained periodic stimulation often induces discomfort, eye strain, fatigue, and drowsiness, and in photosensitive individuals may elevate seizure risk (Fisher et al., 2005; Zhu et al., 2010). These adverse effects restrict session duration, limit participant tolerance, and ultimately affect BCI performance.

To mitigate discomfort while preserving performance, several approaches have focused on stimulus design. Some broaden the usable flicker-frequency range (Bąsaklar et al., 2019; Chabuda et al., 2019; Lai et al., 2024), while others reduce contrast or luminance modulation depth (Cabrera Castillos et al., 2023; Ladouce et al., 2022; Martınez-Cagigal et al., 2023). For example, near-threshold contrast modulations can reliably tag spatial attention (Ladouce & Dehais, 2024), and subtle 14 Hz overlays can track group-level attentional fluctuations during monitoring tasks (Ladouce et al., 2025). Beyond amplitude manipulations, textured visual patterns—flickering stimuli with spatial structure such as orientation, edges, or fine-grained contrast variations rather than uniform luminance—have been explored for their effects on comfort and neural responses (Meng, Liu, et al., 2023; Ming et al., 2023; Schrag et al., 2025). However, these textures can still be strongly distracting in peripheral vision when multiple stimuli are presented simultaneously.

To address this, some authors (Ployart et al., 2022) introduced subtly textured, near-invisible flickers composed of small, randomly oriented elements, engineered to engage pathways from contrast-sensitive retinal ganglion cells to orientation-selective cortical neurons. A key advantage of these stimuli is their low peripheral salience—rods are relatively insensitive to contrast and mainly signal luminance changes—thereby limiting distraction when multiple flickers are presented concurrently. Building on this idea, we replaced these elements with high spatial-frequency Gabor patches (Dehais et al., 2024). When paired with brief aperiodic bursts in a code-modulated VEP (c-VEP) scheme, these Gabor-based textures evoke robust cortical responses and increase event-related potential (ERP) amplitudes by up to a factor of three compared to traditional black-white flicker, enabling accurate single-trial detection with dry-electrode EEG and requiring under one minute of calibration in reactive BCI settings. Yet, these evaluations rely on low-rate, aperiodic presentations (≈ 3 flashes/s), leaving open whether such textured stimuli remain effective in strictly frequency-defined paradigms such as SSVEP.

In the present study, we test whether texture-based stimulation can be integrated into the SSVEP regime while preserving visual comfort and enhancing neural responses, thereby opening new methodological avenues for both cognitive neuroscience and BCI. Previous work has shown that texture can modulate both perceived comfort and the amplitude of elicited SSVEPs (Schrag et al., 2025), but these effects have not been characterized systematically across frequencies, despite the strong frequency dependence of SSVEP responses and the likelihood that texture-related effects also vary with frequency. Importantly, improvements observed for centrally viewed stimuli do not necessarily reduce distraction caused by neighboring peripheral flickers. Our approach comprises two steps. (i) Using a frequency-sweep design (3-18 Hz), we compare traditional plain flickers with Gabor-based textured flickers to identify the frequencies that maximize SSVEP amplitude and to characterize how texture affects neural entrainment and perceptual experience. (ii) Based on these findings, we design a proof-of-concept offline five-class BCI using the optimal frequency range for each stimulus type, assessing multi-class feasibility through classification accuracy and subjective ratings. By jointly analyzing objective neural responses and subjective evaluations of usability, we directly quantify the performance-comfort trade-off and demonstrate how stimulus design shapes both the robustness of frequency-locked activity and the perceptual cost of the paradigm. This dual perspective advances SSVEP-based BCIs toward stimulus designs that combine high reliability with sustained comfort for real-world use outside the laboratory.

## 2 Methods

A video detailing the experimental protocol is available at the following link : https://youtu.be/EvOLoxqT8ss

### 2.1 Participants

Twenty-four healthy participants (7 women, mean age = 27 ± 7 years) took part in the first experimental session, primarily staff and students at ISAE-Supaero. Of these, twenty-one participants returned two months later for the second session (the remaining three could not be reached for scheduling) and completed the second session (7 women, mean age = 27 ± 6 years). For this second session, three participants were excluded from the analysis due to poor EEG signal quality or signs of drowsiness during the experiment, resulting in eighteen participants for the analysis.

All participants provided written informed consents prior to the experiment and were compensated €20 for completing both sessions. The research received ethical approval from the University of Toulouse (CER approval number 2025 1002) and complied with the Declaration of Helsinki. Exclusion criteria included neurological disorders or psychoactive medication use at the time of the study. All participants had normal or corrected-to-normal vision.

Participants were free to take breaks between trials for as long as needed and were encouraged to rest if they experienced any discomfort. They were instructed to inform the experimenter immediately in case of headache, excessive eye strain, or visual fatigue.

### 2.2 EEG recordings

For both sessions, participants were equipped with an 8-channel semi-dry, wireless EEG system (Smarting Pro, mBrainTrain), sampled at 500 Hz. The 8 electrodes were placed according to the 10-20 system over the occipital and parieto-occipital cortex (PO7, O1, Oz, O2, PO8, PO3, POz, PO4) to efficiently capture visually evoked potentials (VEP) and reduce dimensionality. The ground and reference electrodes were positioned at AFz and FCz, respectively. Electrode impedance was maintained below 40kΩ during setup. Recordings were conducted in a standard experimental room, without the use of a Faraday cage. EEG data were recorded using the Lab Streaming Layer (LSL) (Kothe et al., 2025).

### 2.3 Stimuli

Visual stimuli were presented using the PsychoPy software package (Peirce, 2007), and displayed on a Dell P2419HC monitor (1920 × 1080 pixels, 60 FPS, peak luminance: 250 cd/m^2^). All stimuli were circular and had a fixed radius of 150 pixels. Background luminance and stimulus size remained constant throughout the experiment. Two types of stimuli were used, as described in Figure 1: *plain* stimuli and textured *Gabor* stimuli.

**Figure 1:**
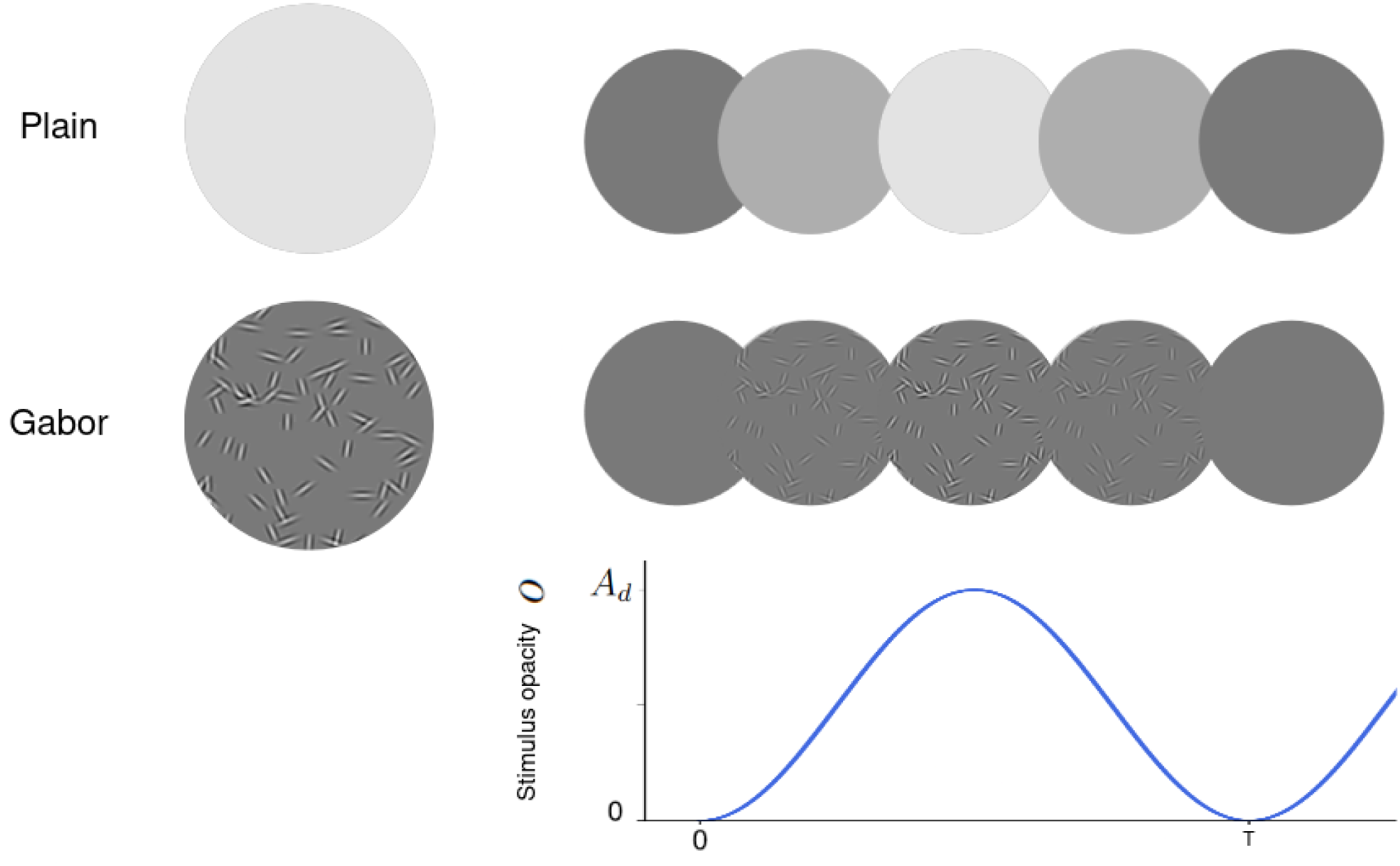
Stimuli and modulation principle used in the experiment. Two types of visual stimuli were used: plain (uniform disks) and Gabor (textured disks composed of sinusoidal gratings). Each stimulus was periodically modulated in opacity following a sinusoidal envelope at a fixed frequency. The right panels illustrate successive frames of the modulation cycle, showing the change in opacity over time. The bottom graph depicts the temporal modulation of the stimulus opacity *o* as a function of time, with *A_d_* representing the modulation amplitude depth and *T* = 1/*f* the period of stimulation.

**Plain stimuli** consisted of a uniform gray disk whose contrast was modulated around a neutral gray background.

**Gabor stimuli** were composed of Gabor patches, following the approach described in Dehais et al., 2024. Each patch consisted of a vertically oriented Gabor filter, characterized by a central white stripe flanked by two black stripes, effectively forming a localized spatial grating. To examine the effect of visual texture on SSVEP responses, we selected a single Gabor stimulus composed of 75 randomly oriented Gabor patches, distributed within the same circular aperture as the plain stimuli, in front of the same neutral gray background. In the following work, Gabor stimuli was kept the same for all participants and all trials, as presented in Figure 1.

To generate SSVEPs, the opacity of each stimulus was sinusoidally modulated over time. To ensure temporal precision and optimize performance, the waveform values were precomputed and stored in an array *W* prior to the experiment. The length of *W* corresponded to the number of display frames per trial, i.e., ⌊*d* · FPS⌋, where *d* is the stimulation duration in seconds and FPS is the frame rate of the monitor. For each frame *i* ∈ [0, ⌊*d* · FPS⌋], the *i*-th element of the waveform and the corresponding stimulus opacity *o*(*i*) were defined as:

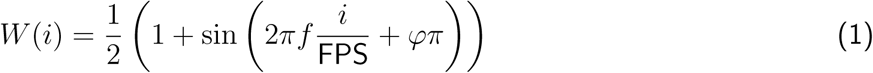

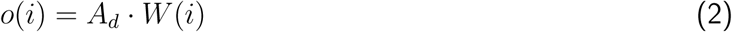

where *f* is the stimulation frequency, *φ* is the phase offset (in half-cycles), and *A_d_* ∈ [0, 1] is the amplitude depth, controlling maximum opacity. This modulation principle and the stimuli are detailed on Figure 1.

### 2.4 Session 1: Frequency Sweep

The first session aimed to assess brain responses to different stimulation frequencies, comparing responses to textured Gabor and plain visual stimuli.

#### 2.4.1 Presentation of the stimuli

All visual stimulation consisted of a single circular stimuli at the center of the screen, either Gabor or plain stimuli. The amplitude depth of the stimulation was set to 100% ranging the contrast from medium gray to white.

Stimulation frequencies spanned from 3 Hz to 18 Hz. Frequencies from 3 Hz to 12 Hz were tested in 1 Hz increments, while frequencies above 12 Hz were sampled every 2 Hz (i.e., 14, 16, and 18 Hz), resulting in 13 discrete frequency conditions. This non-uniform sampling was based on pre-experimental pilot tests, which indicated that the majority of SSVEP power and variability occurs below 12 Hz. Reducing the sampling density at higher frequencies allowed for a shorter session duration, improving participant comfort without compromising the spectral resolution in the most informative range. Once acquired, events containing the displayed frequency were sent using LSL after each block of stimulation to be used for data processing.

#### 2.4.2 Procedure

Participants were comfortably seated at a viewing distance of approximately 70 cm from the screen in a dimly lit room. They were instructed to maintain their gaze on the center of the screen and to actively attend to the stimulus during each trial.

Each trial consisted of 12 stimulation blocks, each lasting 2.2 seconds, separated by a 1 second inter-block interval. During each block, a single frequency and stimulus texture (plain or Gabor) was presented. The order of frequencies and stimulus textures was randomized across trials and participants to counterbalance potential fatigue and order effects, by randomly shuffling the frequencies to be displayed.

No fixation cross was displayed during the stimulation period to avoid introducing additional visual elements that could interfere with the perception of the textured or plain stimuli. Instead, a visual cue appeared as a red square surrounding the central circle during the inter-block interval to alert participants to the upcoming stimulation. The trial course is detailed on Figure 2.

**Figure 2:**
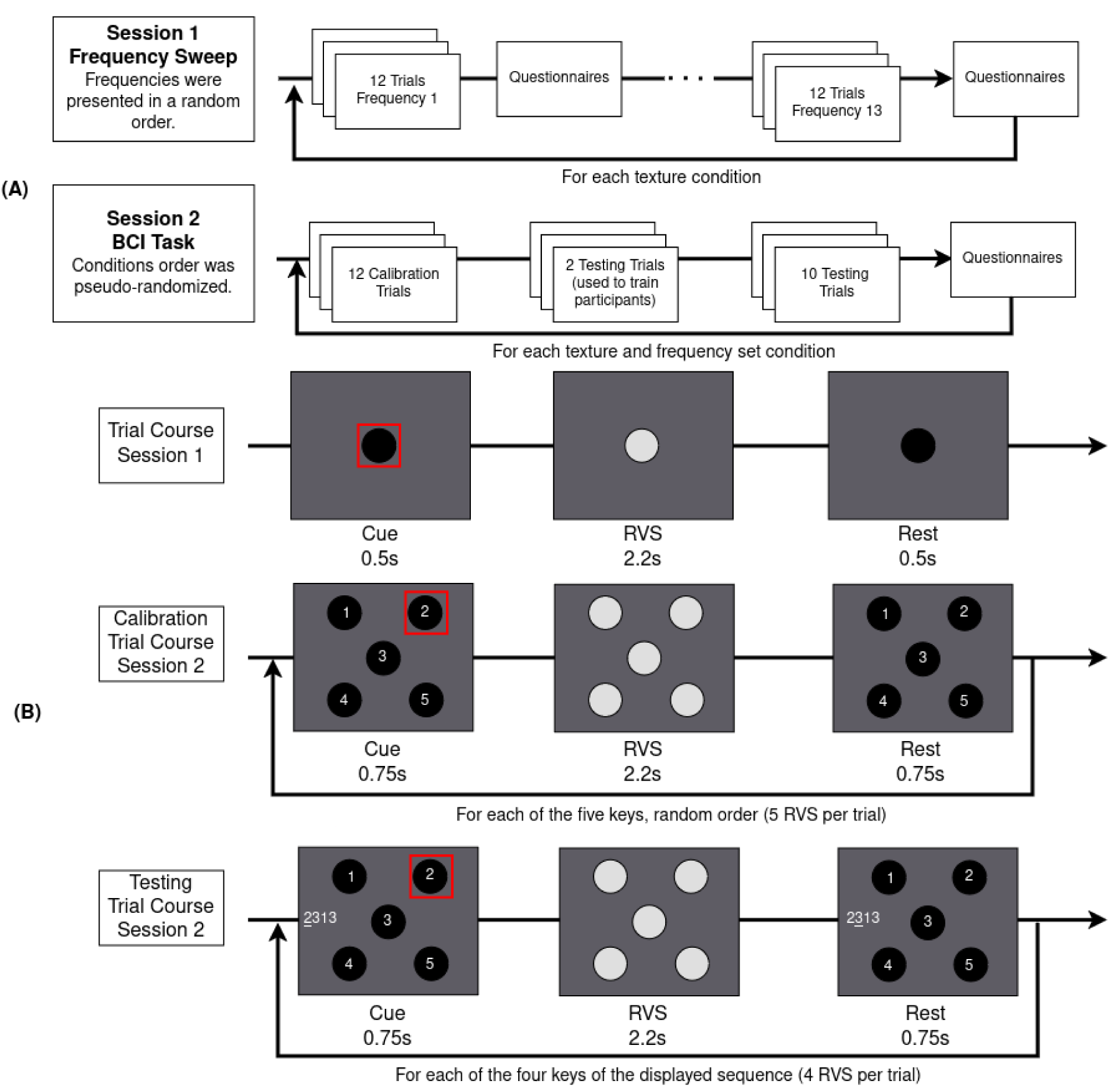
Experimental protocol of the two sessions. **(A)** : Structure of the two sessions. In Session 1, participants underwent a frequency sweep for each texture condition in which each stimulation frequency from the tested set was randomly presented in 12 trials. After each block, participants completed subjective questionnaires. In Session 2, participants performed a BCI task using a 5-key pinpad layout. During each trial, they were cued to fixate a specific target while it flickered at its assigned frequency. Each condition (2 textures × 2 frequency bands) included 24 trials — 12 used for calibration and 12 for testing, including 2 training trials — followed by subjective ratings. **(B)** : trial timeline for each session. In the frequency-sweep, the central target was cued before the onset of the RVS, followed by a resting period. In the BCI calibration trials, one of the five targets was cued before RVS onset, and each complete trial comprised the sequential cueing of all five targets. During the testing trial, a random sequence of 4 digits to be fixated was displayed on the left and digits were cued before RVS onset.

After completing the 12 trials for a given frequency, participants provided subjective ratings on the visual stimulation (see 2.4.4).

The experiment lasted for around 35 minutes including the consent form signature, EEG preparation and visual stimulation recordings.

#### 2.4.3 EEG Processing

EEG data were processed using MNE-Python 1.10.1 (Gramfort et al., 2014) in Python 3.10.

A bandpass filter was applied from 0.5 Hz to 40 Hz to filter low drifts and remove high frequency artifacts while keeping signal of interest (Cohen, 2014). The data were then segmented into epochs corresponding to each stimulation event. For each frequency and stimulus texture (plain vs. Gabor), EEG data were epoched from 0 to 2.2s post-stimulus onset, without baseline correction, resulting in 12 epochs per participant per frequency-stimulus condition.

We did not apply artifact-correction techniques such as Independent Component Analysis (ICA) or Artifact Subspace Reconstruction (ASR) as our objective was to evaluate the stimuli under preprocessing conditions that remain compatible with online BCI deployment. SSVEPs are known to be relatively resilient to typical EEG artifacts (Norcia et al., 2015), and our goal was to keep the pipeline lightweight, reproducible and aligned with practical BCI constraints.

#### 2.4.4 Metrics

Two types of metrics were recorded during this session:

**Objective metrics:** We computed the SSVEP signal-to-noise ratio (SNR) following approach from Cohen and Gulbinaite, 2017. EEG data were first reduced to a single optimized component time series using Rhythmic Entrainment Source Separation (RESS), which enhances frequency-specific activity through spatial filtering. SNR was then defined as the ratio of power at the stimulation frequency to the average power in the surrounding frequency bins within ±1 Hz, excluding the immediate neighboring bins (±0.5 Hz around the target). This approach provides a robust estimate of frequency-specific entrainment by minimizing spectral leakage and avoiding contamination from adjacent peaks.

To investigate the consistency of EEG spectral phase across trial, Inter Trial Coherence (ITC) was considered as an additional metric (Cohen, 2014; Makeig et al., 2002; Tallon-Baudry et al., 1996). ITC at the stimulation frequency was computed separately for each frequency and texture condition using time-frequency decomposition based on complex Morlet wavelets (with compute_tfr(method=“morlet”) method in MNE-Python 1.10.1 following the approach of “Frequency and Time-Frequency Sensor Analysis — MNE 1.11.0 Documentation”, n.d.). For each frequency of stimulation, a single-frequency decomposition was performed at the frequency of stimulation, and ITC was extracted directly from the complex wavelet coefficients returned by the transform. This procedure yielded one ITC time course per channel, which was then averaged over the duration of the stimulation period and across the eight electrodes, resulting in a single ITC value per condition and participant that reflects the consistency of phase across trials at the stimulation frequency.

**Subjective metrics:** After each block, participants rated their experience on an 11-point Likert scale (0-10) for the following dimensions: Visual comfort (0 = very uncomfortable, 10 = very comfortable) and Mental fatigue (0 = none, 10 = extreme), defined as the subjective feeling of cognitive tiredness emerging from sustained attention. Ratings were collected using an on-screen form implemented in PsychoPy, presented after each trial. This resulted in one subjective rating per combination of stimulation frequency and stimulus texture.

### 2.5 Session 2: 5-class BCI task

The second session was designed to evaluate the usability and performance of textured stimuli in an applied BCI context. To this end, participants performed a 5-class SSVEP typing task using the frequency ranges previously identified as optimal for Gabor and Plain stimuli during the first session.

#### 2.5.1 Presentation of the stimuli

In this session, five stimuli were presented simultaneously, spatially arranged around the screen center in a cross-like configuration, as shown in Figure 2, on the bottom panel. Their center coordinates in pixels, relative to the screen center were as follows: [−350, 350], [350, 350], [0, 0], [−350, −350], [350, −350]. This arrangement positioned one stimulus at fixation and four others diagonally around it. Each stimulus was labeled with a number from 1 to 5.

Each stimulus flickered at a distinct frequency, thereby defining five separate SSVEP classes. The exact stimulation frequencies were determined from Session 1: the two most effective frequency ranges based on SSVEP entrainment and subjective comfort were extracted separately for plain and textured stimuli from results of Session 1 (see 3.1.3 for quantitative details). As this session aimed to evaluate performance in a more applied BCI control task than the frequency sweep, the modulation depth was set to 80% for plain stimulation to improve visual comfort for participants (Ladouce et al., 2022).

#### 2.5.2 Procedure

Participants were comfortably seated at approximately 70 cm from the display screen. The order of the four experimental conditions (stimulus texture × frequency set) was pseudo-randomized across participants using a latin square design to counterbalance order effects.

Each condition began with a calibration phase consisting of 12 repetitions per class, i.e., 60 stimulation blocks (5 classes × 12 repetitions). Each block lasted 2.2 s and was followed by a 1.5 s inter-trial interval (ITI). Compared to Session 1, this longer ITI gave participants time to identify the upcoming target location and helped minimize eye movements during visual stimulation.

After the presentation of the calibration trials, participants completed 2 testing trials to familiarize themselves with the task excluded of the analysis, followed by 10 testing trials. During each of these trials, a sequence of 4 digits (from 1 to 5) was displayed on the left side of the screen. Each digit corresponded to one of the five flickering targets to be attended at. Participants were instructed to select the sequence by directing their gaze to the corresponding stimulus, one at a time, resulting in 48 stimulation blocks. Visual cues were still displayed before each digit to indicate the required target. This approach for the task allowed for transferable results to an online classification task. Taking into account 12 trials for calibration, 2 testing trials used to train the participant and 10 testing trials, each condition resulted in 24 trials as described in Figure 2 and thus in 108 stimulation blocks, including 8 blocks corresponding to training excluded of the analysis.

In this session, participants performed the BCI task under four conditions, crossing the two stimulus types (plain vs. Gabor) with the two frequency sets (plain-optimized vs. Gabor-optimized).

After completing each condition, participants provided subjective ratings using three 11-point Likert scales, assessing visual comfort of the stimulation, mental fatigue of the task and stimulation and peripheral distraction from surrounding stimuli.

#### 2.5.3 EEG processing and classification

EEG data were bandpass-filtered between 0.5 and 40 Hz, a frequency range commonly used to preserve steady-state visual evoked potentials while attenuating slow drifts and high-frequency noise such as muscular activity or line noise (Cohen, 2014). The filtered signals were then segmented into epochs of 2.2 s, matching the stimulation duration, with the start of each epoch shifted by a delay offset *d* relative to stimulus onset. This offset, fixed at 135 ms based on pre-test classification results, was introduced to minimize contamination from early visual transients and optimize the extraction of steady-state responses. Epochs corresponding to the training trials were excluded of the analysis.

For offline classification, we implemented a task-related component analysis (TRCA) model (Nakanishi et al., 2018), which maximizes reproducibility of SSVEP responses across trials of the same class. For classification, the TRCA model was fitted exclusively on the calibration trials and its performance assessed on the testing trials.

TRCA was implemented in Python using implementation from python-meegkit 0.17 (Barascud et al., 2023) and the MNE-Python ecosystem (Gramfort et al., 2014).

#### 2.5.4 Metrics

The main performance metric was classification accuracy, defined as the percentage of correctly identified targets in the testing phase, computed per participant.

Subjective metrics (visual comfort, fatigue and peripheral distraction) were analyzed in parallel to provide a comprehensive evaluation of BCI usability during the task using 11-point Likert scales.

### 2.6 Statistical Analyses

For Session 1, objective performance metrics (SNR, ITC) and subjective ratings were analyzed using non-parametric Wilcoxon signed-rank tests for pairwise comparisons at fixed frequencies, as normality assumption was violated in most combination of frequency and texture, as disclosed by Shapiro-Wilk test. Multiple frequency-wise comparisons were corrected using Benjamini-Hochberg False Discovery Rate (FDR) correction, which is appropriate for multiple correlated frequency-wise tests and offers a good compromise between Type-I error control and sensitivity. Effect sizes were reported as matched-pairs Rank-Biserial Correlation (*RBC*) and Common Language Effect Size (*CLES*).

For Session 2, classification accuracies were compared across texture conditions (plain vs. Gabor) separately for each frequency set (Gabor optimized vs. Plain optimized). Wilcoxon signed-rank tests were used as normality assumption was again violated, as disclosed by Shapiro-Wilk test. As only two conditions were tested within each frequency set, no correction for multiple comparisons was applied. Effect sizes (*RBC*, *CLES*) were reported for all tests.

Statistical analyses were conducted in Python using the Pingouin package (Vallat, 2018).

## 3 Results

### 3.1 Results of session 1 : Frequency Sweep

The aim of Session 1 was to systematically examine the influence of stimulus texture and stimulation frequency on SSVEP responses. By studying objective (SNR, ITC) and subjective (comfort, fatigue) metrics across a broad frequency sweep, we sought to identify whether textured stimuli offer neural or ergonomic advantages over classical flickers.

#### 3.1.1 Objective metrics

Figure 3 shows the evolution of SSVEP signal-to-noise ratio (SNR) across stimulation frequencies for plain and Gabor stimuli. Visual inspection suggests a clear frequency-dependent modulation of entrainment strength, with Gabor textures eliciting stronger responses in the low-frequency range, and plain stimuli becoming more effective at higher frequencies. These observations were statistically confirmed by Wilcoxon signed-rank tests conducted at each frequency (Table A). Significant differences were found at 3–9 Hz, where Gabor stimuli consistently produced higher SNR values (*p_corr_ <* 0.05), In contrast, at higher frequencies (10,12,14, 16, and 18 Hz), plain stimuli outperformed Gabor (*p_corr_ <* 0.05). No significant differences were observed at 11Hz.

**Figure 3:**
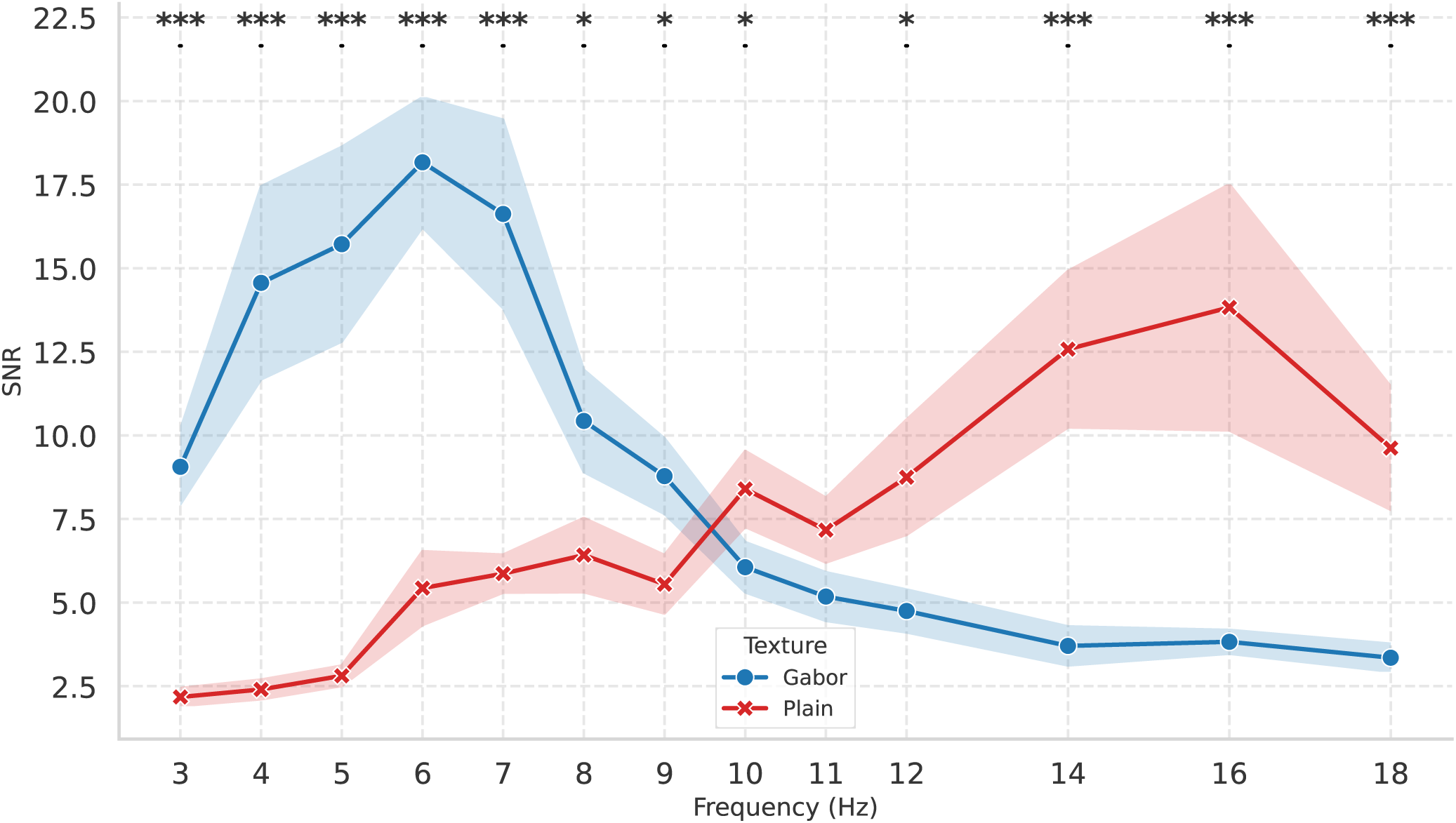
SSVEP signal-to-noise ratio (SNR) as a function of stimulation frequency for plain and Gabor stimuli. Each curve represents the mean across participants (*N* = 24), with shaded areas denoting the standard error of the mean (SEM). The figure illustrates the frequency-dependent entrainment strength and highlights differences between stimulus textures. Asterisks indicate significant differences between Gabor and Plain stimuli at each fixed frequency (Wilcoxon signed-rank tests, FDR-corrected. * : *p <* 0.05, ** : *p <* 0.01, *** : *p <* 0.001). Statistical analysis is detailed in appendix Table A.

Figure 4 shows the evolution of SSVEP ITC at the stimulation frequency averaged across all the electrodes as a function of stimulation frequency for plain and Gabor stimuli. In a similar way as with SNR, ITC shown a strong frequency crossover effects with low frequencies (*<*10 Hz) resulting in higher ITC with Gabor stimuli compared to plain stimuli, and the opposite tendancy in higher frequencies (*>*10 Hz). These observations were statistically confirmed by Wilcoxon signed-rank tests conducted at each frequency (Table A). Significant differences were found on the 3–9 Hz range, where Gabor stimuli consistently resulted in higher ITC values (RBC *>* 0.85, CLES *>* 0.79, *p* < .05),. In contrast, at higher frequencies from 12 Hz to 18 Hz, plain stimuli outperformed Gabor, as reflected by negative RBC values of large magnitude (|*RBC*| *>* 0.80, CLES *>* 0.80, *p* < .05). No significant differences were observed at 10 Hz and 11 Hz.

**Figure 4:**
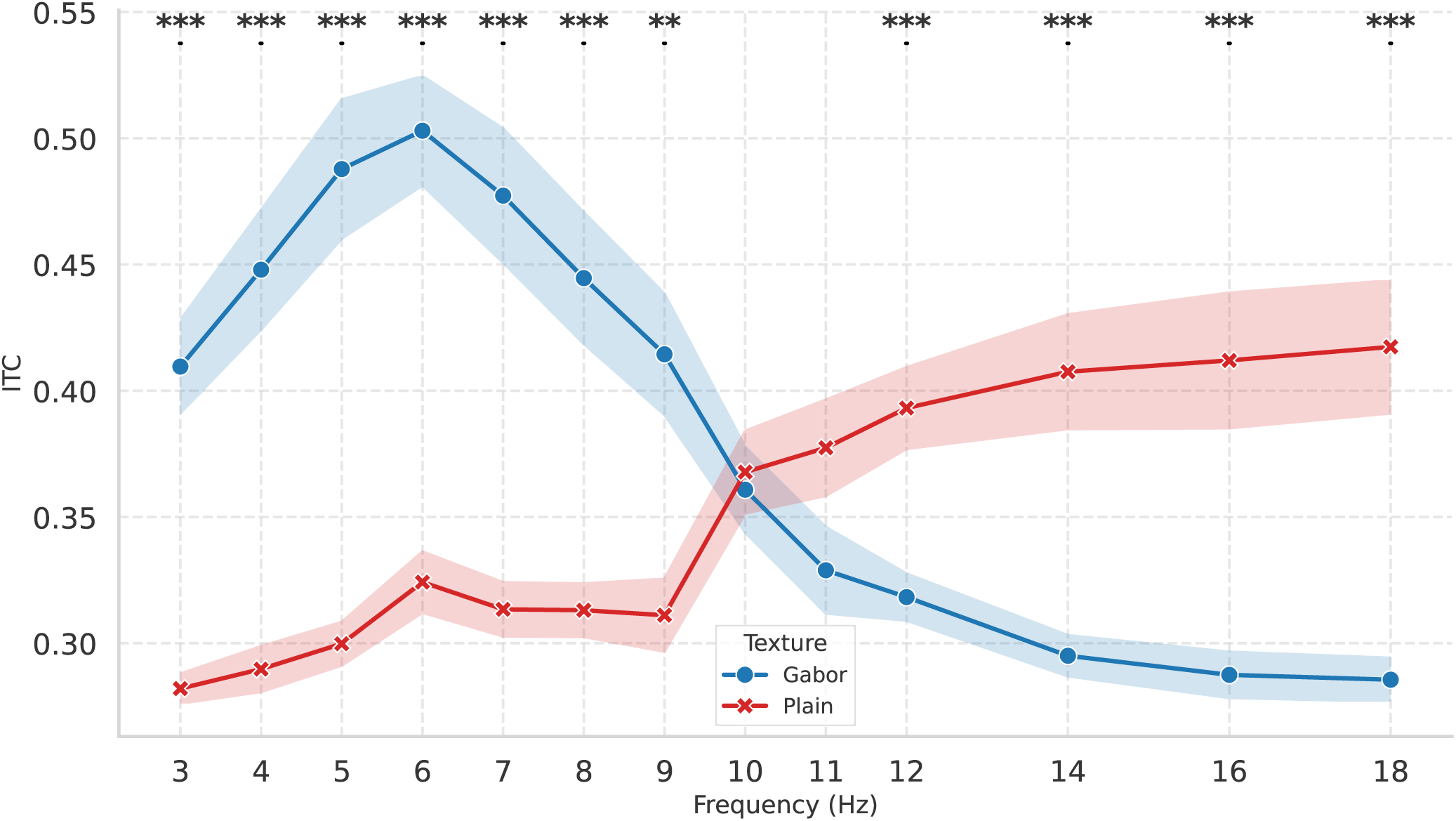
SSVEP inter trial coherence (ITC) at the stimulation frequency averaged across electrodes as a function of stimulation frequency for plain and Gabor stimuli. Each curve represents the mean across participants (*N* = 24), with shaded areas denoting the standard error of the mean (SEM). The figure illustrates the frequency-dependent entrainment strength and highlights differences between stimulus textures. Asterisks indicate significant differences between Gabor and Plain stimuli at each fixed frequency (Wilcoxon signed-rank tests, FDR-corrected. * : *p <* 0.05, ** : *p <* 0.01, *** : *p <* 0.001). Statistical analysis is detailed in supplementary Table 2.

Together, these results indicate a robust crossover effect: Gabor textures are more effective in driving SSVEP responses at lower frequencies, while plain flickers dominate at higher stimulation rates.

#### 3.1.2 Subjective metrics

Having established the objective SSVEP profile, we complemented the analysis with subjective metrics to evaluate user-centered aspects of the stimulation experience.

Subjective ratings showed a consistent preference for textured Gabor stimuli over plain flickers (Figure 5).

**Figure 5:**
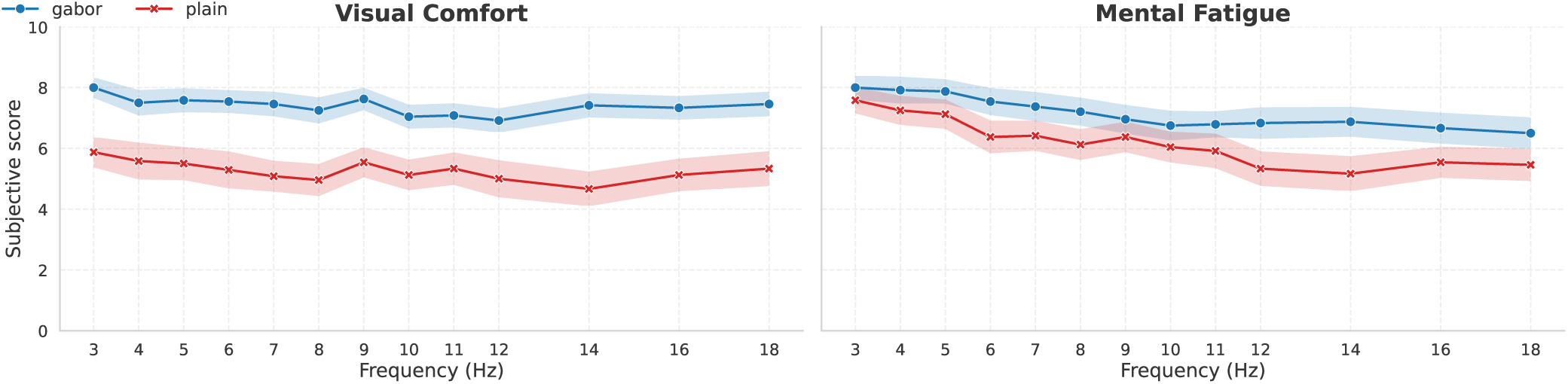
Subjective ratings of visual comfort and mental fatigue as a function of stimulation frequency. Ratings were provided on an 11-point Likert scale (0-10) in such a way that the higher the rating, the better. Each curve represents the mean across participants (*N* = 24), with shaded areas denoting the standard error of the mean (SEM) for plain and Gabor stimuli.

For *visual comfort*, Gabor stimuli were rated as more comfortable across all tested frequencies, with Wilcoxon signed-rank tests confirming effects at all frequencies (*p*_corr_ < .05) except 4Hz and 12Hz.

By contrast, *mental fatigue* was less consistently modulated by stimulus type, with only a single significant effect observed at 14 Hz confirmed by Wilcoxon test (*p*_corr_ < .05).

Significant statistical analyses results are summarized in appendix Table 3.

These results indicate that beyond their objective advantages at low stimulation frequencies, textured flickers substantially improve subjective user experience, particularly by increasing comfort.

#### 3.1.3 Selection of Frequency Sets for Session 2

The frequency sweep conducted in Session 1 revealed a clear crossover pattern across textures. Gabor stimuli elicited significantly stronger SSVEP responses at lower stimulation frequencies (3-9 Hz; see Table A), whereas plain flickers became more effective in the higher range (12-18 Hz).

Based on this combined evidence, two optimized frequency sets were defined for the multi-class BCI task in Session 2. The *Gabor-optimized* set (5-7 Hz) corresponded to the band yielding the strongest frequency-specific entrainment and most favorable subjective evaluations for textured stimuli. The *Plain-optimized* set (14-16 Hz) captured the range where plain flickers consistently outperformed Gabor textures in terms of SSVEP amplitude.

These two sets formed the basis of the four experimental conditions in Session 2 (crossing texture × frequency set).

### 3.2 Results of session 2 : 5-class BCI task

The second session focused on assessing both objective and subjective measurements during a 5-class BCI task comparing Gabor and Plain stimuli. This task used two distinct frequency sets (optimized for either Gabor or Plain stimuli) derived from the first session’s command encoding (Part 3.1.3).

Consequently, the Latin square counterbalancing resulted in an uneven distribution of participants across the four condition pairs (five participants for two pairs, and four for the other two). To ensure that this imbalance did not introduce systematic bias, we tested for potential order effects on both objective and subjective measures. Since data distributions violated normality assumptions, non-parametric Kruskal-Wallis tests were applied. The order of condition presentation had no significant effect on BCI accuracy (*H* = 3.78, *p* = 0.29), visual comfort ratings (*H* = 1.12, *p* = 0.77), mental fatigue ratings (*H* = 1.69, *p* = 0.64), or reported peripheral distraction (*H* = 1.39, *p* = 0.71). Collectively, these results indicate that the minor unbalance in the Latin square design did not bias the outcomes. Therefore, all subsequent analyses were conducted on the 18 remaining participants.

#### 3.2.1 Objective metrics

On average, for the Gabor-optimized band, classification accuracy was higher with Gabor stimuli (*M* = 95.7%, *SD* = 14.2%) than with Plain stimuli (*M* = 79.9%, *SD* = 18.7%). A two-sided Wilcoxon signed-rank test confirmed that this difference was statistically significant, *W* = 0.0, *p* = 0.002, *RBC* = 1, *CLES* = 0.81.

For the plain-optimized band, classification accuracy was higher with Plain stimuli (*M* = 98.1%, *SD* = 4.7%) compared to Gabor stimuli (*M* = 78.6%, *SD* = 26.4%). This difference was also statistically significant, *W* = 2.5, *p* = 0.001, *RBC* = 0.963, *CLES* = 0.835.

Overall, both stimulus types yielded their strongest classification performances within their respective optimal frequency bands. Beyond these objective results, we also evaluated participants’ subjective experience to assess the ergonomic impact of the stimulus design.

#### 3.2.2 Subjective metrics

In addition to BCI accuracy, we studied subjective ratings on 11-point Likert scales regarding visual comfort, mental fatigue and peripheral distraction during the task.

##### Visual comfort

Gabor stimuli (*M* = 7.4, *SD* = 1.6) were noted more visually comfortable than plain stimuli (*M* = 4.3, *SD* = 2.2) for the Gabor-optimized band. A two-sided Wilcoxon test confirmed that this different was statistically significant, *W* = 10, *p* = 0.0051, *RBC* = 0.87, *CLES* = 0.86.

For the plain-optimized band, Gabor stimuli (*M* = 7.3, *SD* = 1.5) were noted more visually comfortable than plain stimuli (*M* = 3.6, *SD* = 2.5). A two-sided Wilcoxon test disclosed that this different was statistically significant, *W* = 5.0, *p* = 0.0051, *RBC* = 0.93, *CLES* = 0.88.

##### Mental Fatigue

Gabor stimuli (*M* = 6.3, *SD* = 2.2) were noted less mental fatiguing than plain stimuli (*M* = 4.4, *SD* = 2.1) for the Gabor-optimized band. A two-sided Wilcoxon test confirmed that this difference was statistically significant, *W* = 24.5, *p* = 0.016, *RBC* = 0.68, *CLES* = 0.75.

For the plain-optimized band, Gabor stimuli (*M* = 6.4, *SD* = 2.4) were noted less mental fatiguing than plain stimuli (*M* = 4.8, *SD* = 2.8). A two-sided Wilcoxon test confirmed that this difference was not statistically significant *p >* 0.05.

##### Peripheral distraction

Gabor stimuli (*M* = 7.4, *SD* = 2.2) were noted less distracting in peripheral regions than plain stimuli (*M* = 4.7, *SD* = 3.1) for the Gabor-optimized band. A two-sided Wilcoxon test confirmed that this difference was statistically significant, *W* = 13.5, *p* = 0.0051, *RBC* = 0.80, *CLES* = 0.74.

For the plain-optimized band, Gabor stimuli (*M* = 7.4, *SD* = 2.5) were less distracting in peripheral regions than plain stimuli (*M* = 3.9, *SD* = 2.9). A two-sided Wilcoxon test confirmed that this difference was statistically significant, *W* = 8.5, *p* = 0.0073, *RBC* = 0.86, *CLES* = 0.82.

## 4 Discussion

The primary goal of this study was to determine whether subtly textured, Gabor-based stimuli can improve both user experience and neural entrainment the SSVEP regime, and whether these benefits translate into accurate multi-class BCI control. To this end, we first carried out a frequency sweep to chart SNR as a function of frequency for texture versus plain luminance flickers, and then tested a five-class BCI using frequency sets optimized from the sweep. Beyond classical objective metrics (SNR and ITC in Session 1; accuracy in Session 2), we collected subjective measures of visual comfort and mental fatigue to quantify end-user experience and relate it to neural performance.

Before examining the neural mechanisms underlying the crossover in SSVEP responses, we first consider participants’ subjective experience of the flicker stimuli, as comfort and perceptual load directly shape the practical usability of SSVEP paradigms.

### 4.1 Gabor stimuli as a tool to improve comfort

Across the full sweep, participants consistently reported higher visual comfort for textured Gabor flickers compared to plain luminance flickers. These effects were further complemented in the multi-target BCI configuration of Session 2, during which participants still reported higher visual comfort and less peripheral distraction for textured Gabor flickers compared to plain luminance stimuli for all frequencies. When several stimuli flicker simultaneously, plain luminance modulation that is tolerable in isolation quickly becomes perceptually dominant, producing strong impressions of flicker and motion and drawing attention away from the attended target (Forschack et al., 2022).

By contrast, Gabor-based textured stimuli attenuate uniform luminance fluctuations and instead drive orientation-selective processing near fixation, thereby reducing peripheral salience and limiting inter-stimulus interference. This behavior is consistent with underlying retinal circuitry: Gabor textures engage contrast-sensitive pathways supported by ON-OFF bipolar cells, which respond to both increments and decrements in luminance and effectively encode edges and spatial transitions—akin to an edge-enhancement filter (Euler et al., 2014; Ichinose & Habib, 2022; Kartsaki, 2022). Because this modulation is spatially structured rather than global, it is predominantly processed foveally, where cone-rich regions of the retina are highly sensitive to contrast and support fine-grained spatial resolution (Jonas et al., 1992). In the periphery, where rod-dominated photoreception favors luminance changes but is less responsive to contrast, the Gabor patches appear weaker and less defined, producing faint texture rather than salient flicker. The densely tiled microstructure of the Gabor stimuli further amplifies this effect through perceptual crowding, whereby many similar elements reduce one another’s recognizability in peripheral vision (Harrison & Bex, 2015).

These perceptual characteristics explain why textured stimulation remained comfortable even with five concurrent targets. This demonstrates that the comfort advantage observed during single-target flicker transfers robustly to realistic multi-target SSVEP layouts. The findings also align with our previous c-VEP study using similar Gabor textures, where participants reported high central comfort and limited peripheral distraction despite repetitive stimulation (Dehais et al., 2024). Together, the results indicate that Gabor-based textures generalize across c-VEP and SSVEP paradigms and offer a practical route to comfortable, high-performance multi-target BCIs as well as for fundamental cognitive neuroscience studies.

While these perceptual differences already point toward functional advantages for textured stimulation, the frequency sweep provides additional insight into how these comfort benefits relate to the underlying neural dynamics of entrainment. We next turn to objective electrophysiological measures to characterize the frequency-dependent crossover between texture and plain flicker.

### 4.2 Two frequency sweet spots and their likely origins

The frequency sweep revealed a robust crossover in the SNR of the SSVEP responses (Figure 3): textured Gabor stimuli outperformed plain stimuli at low frequencies (3-9 Hz), while plain stimuli excelled at higher frequencies (12-18 Hz). Importantly, this low-frequency advantage for texture was not only reflected in amplitude but also in phase consistency across trials (Figure 4). Phase coherence was markedly higher for texture in the 3-9 Hz range, indicating that there was less variability in the timing of SSVEP using Gabor stimuli at low frequency compared to plain luminance stimuli. At higher frequencies (12-18Hz), phase coherence of Gabor-elicited responses diminished in favor of plain stimuli. This motivated two application-oriented “sweet spots”, later validated in the multi-class BCI task: a low-frequency band optimized for textured stimuli and a higher-frequency band optimized for plain stimuli.

The high-rate advantage for plain flicker aligns with classic SSVEP tuning to alpha-beta luminance modulation (e.g., Ladouce et al., 2022; Nakanishi et al., 2018; Norcia et al., 2015). On the other hand, the low-rate advantage for texture highlights a complementary regime in which spatial structure yields superior entrainment together with improved comfort. This interpretation accords with our previous study that slow pace aperiodic textured Gabor stimulation elicits significantly higher ERPs amplitude than plain flicker (Dehais et al., 2024). This low-frequency sweet-spot for textured stimuli also matches frequency-tagging work using complex visual content, where faces, words, and other structured stimuli are typically presented in the 4-8 Hz range that best reveals category-sensitive responses (e.g., Boremanse et al., 2013; Gruss et al., 2012; Rossion et al., 2012). Although our textures are not categorical, they share critical spatial features and appear to operate optimally in the same low-frequency window.

A coherent explanation of this crossover begins with considerations of the Critical Flicker Fusion (CFF) threshold, which is the frequency at which flicker is perceived as continuous. Gabor-textured stimuli are intentionally subtle and locally structured. As stimulation frequency increases, their effective temporal contrast decreases more steeply than that of plain luminance flicker. Because CFF strongly depends on both contrast and spatial frequency (Mankowska et al., 2021), Gabor stimuli are expected to reach perceptual fusion earlier. Beyond approximately 12-14 Hz, this reduction in perceived flicker naturally attenuates the resulting steady-state response, consistent with the drop in SSVEP SNR observed in our data.

Beyond perceptual fusion, a second factor concerns how textured and plain flicker could recruit different spatio-temporal mechanisms in the early visual system. The spatial structure of the Gabor stimuli—defined by localized edges, orientations, and high spatial-frequency content—suggests a differential engagement of the two major visual pathways, magnocellular (M) and parvocellular (P) (Masri et al., 2020). These pathways exhibit well-established differences in temporal and spatial tuning: the M pathway is more sensitive to rapid luminance changes and lower spatial frequencies, whereas the P pathway preferentially processes fine spatial details and shows reduced sensitivity to higher temporal rates (Goodhew et al., 2014). Although the exact separation is not absolute, as high spatial frequencies can also elicit M-pathway responses (Skottun, 2015), the textured Gabor flickers likely place a greater relative load on P-dominated mechanisms than plain luminance flicker. This distinction offers a plausible account for the frequency-dependent advantage observed in our results. Because the P pathway exhibits lower temporal sensitivity, P-driven responses should degrade more quickly as flicker frequency increases; conversely, at slower stimulation rates, the richer spatial content of the Gabor mosaic can be more effectively integrated, yielding stronger entrainment. Prior work examining pathway contributions to SSVEPs supports a frequency-dependent separation, although the exact ranges remain debated: F. B. Vialatte et al., 2009 reported P-pathway dominance around ∼16 Hz with M-pathway dominance at higher frequencies, whereas Kristensen et al., 2016 estimated P contributions up to ∼10 Hz and M contributions from ∼10-20 Hz. Despite these discrepancies, both accounts situate our observed Gabor advantage (5-7 Hz) within a range where P-mediated responses are expected to remain robust, while the decline of Gabor SNR at higher rates is consistent with the reduced temporal bandwidth of P mechanisms. Although these interpretations are plausible and consistent with known spatio-temporal tuning of the M and P pathways, they remain indirect and would benefit from dedicated experiments combining SSVEP with contrast-response characterization.

Altogether, these considerations suggest that the richer spatial structure of the Gabor stimuli facilitates more reliable entrainment dynamics, strengthening both neural entrainment and phase stability with sufficient time per cycle, at low frequencies.

### 4.3 Applications of textured stimuli to BCI control

From a BCI perspective, the identified frequency bands point to a clear functional opportunity: in the low-frequency regime, Gabor stimuli consistently yielded higher SNR and ITC, indicating stronger, more stable entrainment. Because these metrics are tightly linked to class separability and decoding reliability, our sweep results demonstrate that textured stimuli provide a promising pathway for robust and comfortable SSVEP-based BCI control. The five-class BCI confirmed the functional value of these frequency-specific sweet spots. Accuracy was high in both bands, with particularly strong performance for texture in the low-frequency condition and minimal inter-participant variability aside from a single outlier. This demonstrates that the benefits observed in the sweep translate directly to practical multi-class control: textured stimuli deliver high classification accuracy while alleviating one of the main drawbacks of SSVEP-based BCIs, namely the salience and visual clutter of multi-target flicker arrays. In other words, low-frequency textured flicker offers a high-performance yet visually unobtrusive command code, making it a compelling option for real-world multi-target BCIs where sustained comfort and peripheral transparency are essential. Textured stimuli approach thus complements recent stimulus-design studies aiming at improving user comfort in BCI, as detailed in the introduction (e.g., Ladouce et al., 2022; Lai et al., 2024; Martınez-Cagigal et al., 2023; Meng, Zhou, et al., 2023; Ming et al., 2023; Schrag et al., 2025), positioning textured stimuli as solid alternative to preserve performance in BCI control with enhanced user comfort.

It is also worth noting that these high accuracies with texture based stimuli, together with improved comfort, were obtained using an 8-channel semi-dry EEG system, which is fast to set up and does not require conductive gel—unlike many SSVEP BCIs that rely on high-density, gel-based caps to maximize SNR. Combined with our texture-based stimulation scheme, this lightweight hardware configuration points toward user-friendly BCIs that can extend beyond the laboratory and into everyday environments.

### 4.4 Limitations and future work

For cognitive neuroscience, textured SSVEPs provide a comfortable and perceptually stable tool for studying neural entrainment, attention, and high-level visual processing with reduced peripheral interference. They complement periliminal and near-threshold tagging approaches that similarly prioritize user tolerance and long-duration feasibility (Ladouce & Dehais, 2024; Ladouce et al., 2025). For BCI research, our results show that textured flicker unlocks a low-frequency operating regime (around 5–7 Hz) that preserves strong neural entrainment while substantially improving visual comfort and reducing peripheral salience, two properties that are crucial for multi-target layouts and prolonged use. This constitutes a qualitatively different strategy from high-frequency or high-contrast designs, and highlights that performance can be maintained even when prioritizing comfort.

Several limitations point to promising directions for future work. Our frequency sweep focused on low and high bands; denser sampling in the 8–12 Hz range would refine the characterization of the crossover. In addition, we tested a single texture recipe. Systematic manipulations of orientation content, spatial frequency, element density, and modulation depth may extend the effective operating range for texture and enable individual optimization. Our BCI evaluation was offline; online, closed-loop studies are needed to assess calibration efficiency, long-session stability, and robustness to eye movements and ecological distractions. Regarding the experimental design, the small sample size (24 participants for session 1 and 18 for session 2) is to be taken into considerations as high variability was found in responses to flickers, as highlighted in SNR (Figure 3), ITC (Figure 4) and accuracy (Figure 6), with the presence of outliers. Finally, our mechanistic interpretation remains indirect: combining SSVEP with source localization, contrast-response measurements, and rapid sweeps across spatial-frequency and orientation dimensions will help disentangle the respective contributions of temporal fusion, feature-selective gain, and attentional modulation.

**Figure 6:**
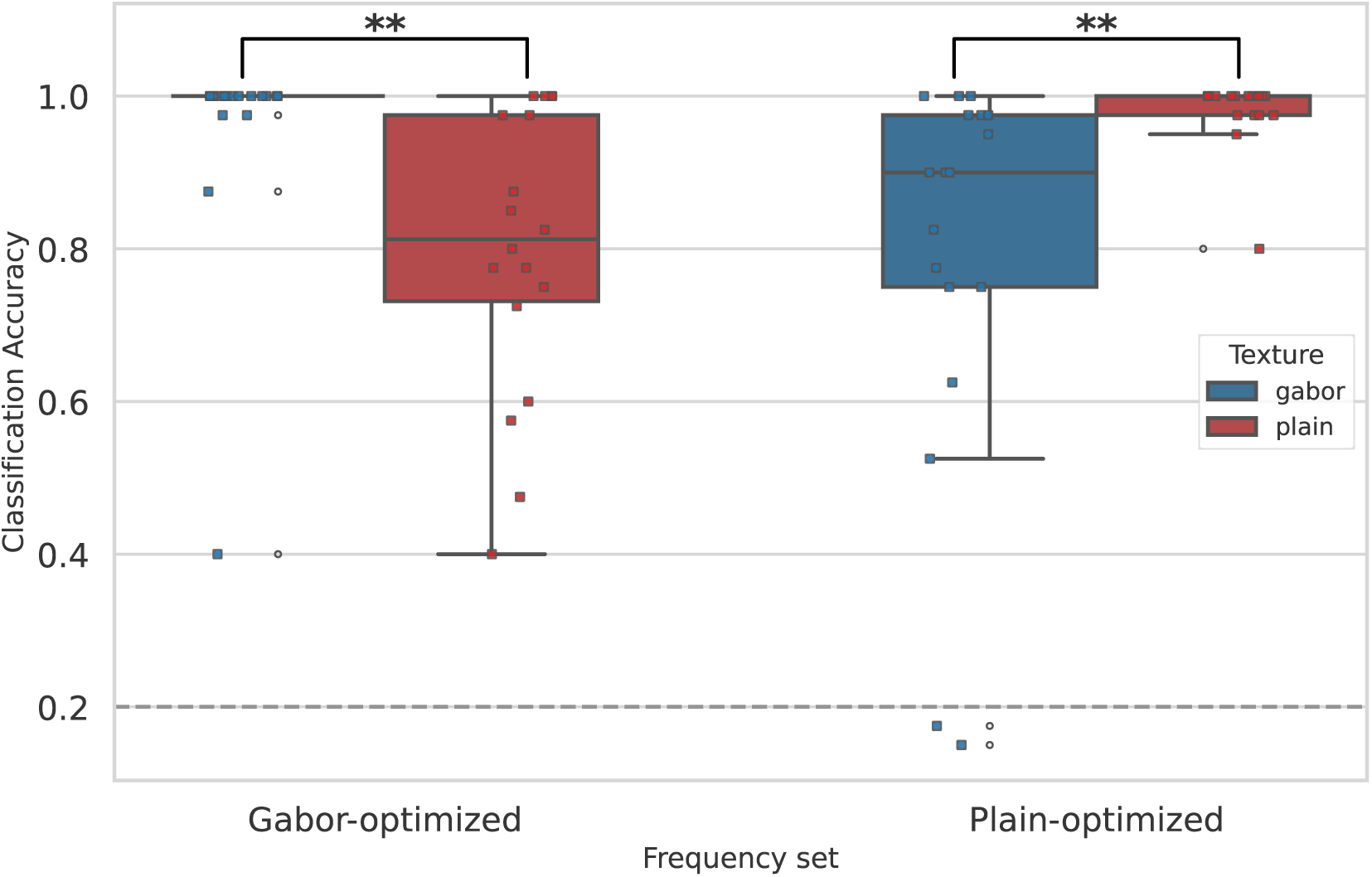
BCI classification accuracy as a function of texture (Gabor or Plain stimuli) and frequency of stimulation (extracted from Session 1, either Gabor-optimized frequencies or Plain-optimized frequencies as described in 3.1.3) across participants (*N* = 18). Asterisks indicate significant differences between Gabor and Plain stimuli at each fixed frequency (Wilcoxon signed-rank tests, FDR-corrected. * : *p* < 0.05, ** : *p* < 0.01, *** : *p* < 0.001).

**Figure 7:**
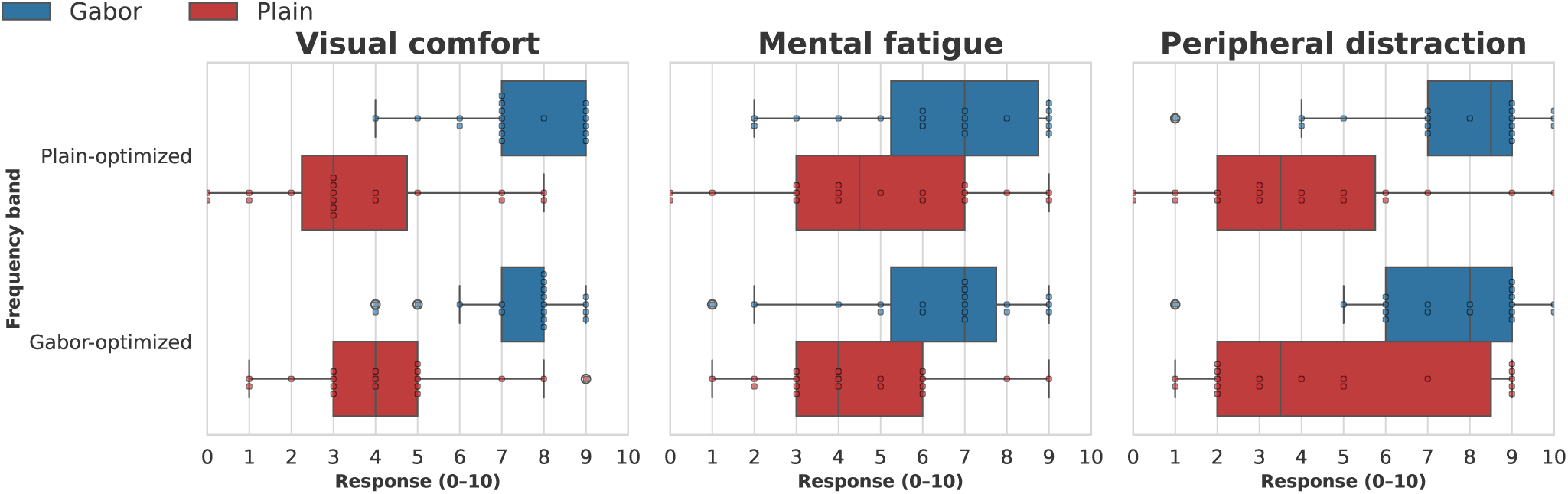
Subjective ratings regarding visual comfort, mental fatigue and peripheral distraction during the stimulation, rated on an 11-point Likert scale as a function of stimulus texture (Gabor and Plain) and the set of frequency used (extracted from Session 1, either Gabor-optimized frequencies or Plain-optimized frequencies as described in 3.1.3) across participants (*N* = 18). Questions were asked in such way that the higher the grade, the better.

## 5 Conclusion

Overall, our findings indicate that textured, Gabor-based flickers provide a comfortable, low-salience alternative to plain luminance flicker. They support strong low-frequency SSVEP responses suitable for multi-class BCI while improving user experience, whereas plain flicker remains advantageous at higher stimulation rates. This complementary pattern yields a principled, frequency-aware stimulus-design space for both cognitive neuroscience—probing cognition and neural entrainment—and practical BCI applications.

Textured Gabor flicker provides a comfortable and perceptually unobtrusive alternative to classical luminance flicker while preserving strong SSVEP entrainment at low frequencies. Across a wide frequency sweep and a multi-class BCI task, Gabor stimuli reliably improved user experience and enhanced phase-locked neural responses in the 3-9 Hz range, whereas plain flicker remained advantageous at higher stimulation rates. This complementary profile reveals a principled, frequency-aware design space for SSVEP paradigms, supporting both cognitive neuroscience studies of neural entrainment and the development of practical, user-friendly BCIs.

Altogether, the present work outlines a pathway toward SSVEP paradigms that are not only efficient but also genuinely user-centered, bridging the gap between experimental entrainment studies and practical, long-duration BCI deployment.

## Data and Code Availability

The dataset is available for non-commercial scientific research, in accordance with participant consent, ethical approval, and GDPR regulations : https://doi.org/10.57745/XVDCWA. The dataset complies to EEG BIDS standard (Pernet et al., 2019).

The code to run the experiment and the analysis is publicly available at the following link : https://doi.org/10.5281/zenodo.17854744.

## Author Contributions

JG : Conceptualization, methodology, software, data collection, analysis, visualization, writing & editing. FD and MCC : Conceptualization, methodology, supervision, validation, writing & editing

## Supplementary Material

A visual abstract summarizing the study and key findings is available at the following link : https://youtu.be/M6C87aY5I-Q

## A Appendix

**Table 1:** Wilcoxon signed-rank test results for SNR in Session 1 (Gabor vs. Plain stimuli). Reported values include Wilcoxon statistic (*W*), corrected *p*-values, rank-biserial correlation (RBC), and common language effect size (CLES). Positive RBC values indicate stronger responses for *Gabor* compared to *Plain*, and negative values indicate the reverse. CLES values indicates the probability of Gabor SNR being higher than plain SNR. Significant *p*-values (*p*_corr_ < .05) are shown in bold.

| Frequency (Hz) | $W$ | $p_{corr}$ | RBC | CLES |
| --- | --- | --- | --- | --- |
| 3 | 10 | **1.7e-05** | 0.93 | 0.91 |
| 4 | 1 | **2.0e-06** | 0.99 | 0.95 |
| 5 | 3 | **3.0e-06** | 0.98 | 0.93 |
| 6 | 24 | **1.7e-04** | 0.84 | 0.91 |
| 7 | 19 | **7.9e-05** | 0.87 | 0.76 |
| 8 | 73 | **0.032** | 0.51 | 0.68 |
| 9 | 62 | **0.015** | 0.59 | 0.69 |
| 10 | 78 | **0.042** | -0.48 | 0.38 |
| 11 | 111 | 0.28 | -0.26 | 0.39 |
| 12 | 69 | **0.025** | -0.54 | 0.32 |
| 14 | 1 | **2.0e-06** | -0.99 | 0.19 |
| 16 | 29 | **3.0e-04** | -0.81 | 0.22 |
| 18 | 18 | **7.8e-05** | -0.88 | 0.16 |

**Table 2:** Wilcoxon signed-rank test results for ITC in Session 1 (Gabor vs. Plain stimuli). Reported values include Wilcoxon statistic (*W*), corrected *p*-values, rank-biserial correlation (RBC), and common language effect size (CLES). Positive RBC values indicate stronger responses for *Gabor* compared to *Plain*, and negative values indicate the reverse. CLES values indicates the probability of Gabor ITC being higher than plain ITC. Significant *p*-values (*p*_corr_ < .05) are shown in bold.

| Frequency (Hz) | $W$ | $p_{corr}$ | RBC | CLES |
| --- | --- | --- | --- | --- |
| 3 | 0.0 | **2.0e-06** | 1.00 | 0.95 |
| 4 | 3.0 | **2.0e-06** | 0.98 | 0.93 |
| 5 | 4.0 | **2.0e-06** | 0.97 | 0.93 |
| 6 | 2.0 | **2.0e-06** | 0.99 | 0.93 |
| 7 | 13.0 | **1.9e-05** | 0.91 | 0.91 |
| 8 | 22.0 | **9.2e-05** | 0.85 | 0.83 |
| 9 | 46.0 | **2.4e-03** | 0.69 | 0.79 |
| 10 | 118.0 | 0.37 | -0.21 | 0.47 |
| 11 | 85.0 | 0.07 | -0.43 | 0.35 |
| 12 | 37.0 | **8.5e-04** | -0.75 | 0.21 |
| 14 | 19.0 | **5.9e-05** | -0.87 | 0.16 |
| 16 | 2.0 | **2.0e-06** | -0.99 | 0.14 |
| 18 | 9.0 | **9.0e-06** | -0.94 | 0.14 |

**Table 3:** Wilcoxon signed-rank test significant results for subjective ratings in Session 1 (Gabor vs. Plain). Positive RBC values indicate higher ratings for Gabor, negative values indicate lower ratings for Gabor. Significant *p*-values after correction (*p*_corr_ < .05) are shown in bold.

| Question | Frequency (Hz) | $W$ | $p_{\text{corr}}$ | RBC | CLES |
| --- | --- | --- | --- | --- | --- |
| Visual comfort | 3 | 22.5 | **0.015** | 0.79 | 0.77 |
|  | 5 | 22.0 | **0.017** | 0.74 | 0.74 |
|  | 6 | 16.0 | **0.015** | 0.81 | 0.73 |
|  | 7 | 6.0 | **0.011** | 0.92 | 0.79 |
|  | 8 | 48.5 | **0.015** | 0.68 | 0.77 |
|  | 9 | 41.5 | **0.017** | 0.67 | 0.78 |
|  | 10 | 43.0 | **0.017** | 0.66 | 0.75 |
|  | 11 | 37.5 | **0.026** | 0.64 | 0.71 |
|  | 14 | 30.0 | **0.006** | 0.80 | 0.79 |
|  | 16 | 43.5 | **0.017** | 0.66 | 0.76 |
|  | 18 | 43.0 | **0.017** | 0.66 | 0.74 |
| Mental fatigue | 14 | 33.0 | **0.017** | -0.69 | 0.32 |

